# Biosorption potential and molecular characterization of metal resistant autochthonous microbes from tannery solid waste

**DOI:** 10.1101/2021.11.12.468455

**Authors:** Hajira Younas, Aisha Nazir, Zakia Latif, Janice E. Thies, Muhammad Shafiq, Firdaus-e-Bareen

**Affiliations:** Institute of Botany, University of the Punjab, Lahore 54590, Pakistan; Institute of Microbiology and Molecular Genetics, University of the Punjab, Lahore 54590, Pakistan; Department of Crop and Soil Science, Cornell University, Ithaca, NY 14853, USA

**Keywords:** *Bacillus*, biosorption potential, heavy metals, tannery solid waste, *Trichoderma*

## Abstract

This study encompasses isolation and screening of heavy metal-resistant fungal and bacterial strains from tannery solid waste (TSW). Twelve fungal strains and twenty-five bacterial strains were isolated from TSW. The growth of fungal strains was observed against different heavy metals ranging from 10 mg L^-1^ to 1050 mg L^-1^ and the growth of bacteria was observed in metal concentrations ranging from 10 mg L^-1^ to 1200 mg L^-1^. Five multi-metal resistant fungal isolates belonging to the genus *Trichoderma* and ten bacterial isolates belonging to the genus *Bacillus* showed good metal resistance and biosorption potential. They were identified through molecular techniques, fungi based on ITS region ribotyping, and bacteria based on 16S rRNA ribotyping. The fungal strains were characterized as *T. hamatum* (TSWF-06), *T. harzianum* (TSWF-11), *T. lixii* (TSWF-02) and *T. pseudokoningii* (TSWF-03, TSWF-10). The bacterial strains were characterized as *Bacillus xiamenensis* (TSW-02), *B. velezensis* (TSW-05), *B. piscis* (TSW-06), *B. safensis* (TSW-10), *B. subtilis* (TSW-14, TSW-15, TSW-17) *B. licheniformis* (TSW-19), *B. cereus* (TSW-20) and *B. thuringiensis* (TSW-22). The fungal strains namely, *T. pseudokoningii* (TSWF-03) and *T. harzianum* proved to be two multi-metal resistant strains with good biosorption efficiency. Unlike fungi, bacterial strains showed metal specific resistance. The strains *Bacillus xiamenensis, B. subtilis* (TSW-14) and *B. subtilis* (TSW-15) showed good biosorption efficiency against Cr, *B. safensis* against Cu, *B. piscis and B. subtilis* (TSW-17) against Pb and *B. licheniformis* and *B. thuringiensis* against Zn. The autochthonous fungal and bacterial strains can therefore be employed to clean metal contaminated environments.

## Introduction

Heavy metals (HMs) pose a serious threat to mankind through increased levels in agricultural lands, water bodies and natural ecosystems. They can be categorized into non-essential metals (As, Cd, Cr, Hg, Ni and Pb) and essential metals (Cu, Fe and Zn) [1]. Various sources of HMs include agricultural activity, industrial effluents, fertilizers, mining and solid waste dumping sites as well as atmospheric sources. Toxic metals have no role in biological pathways and their excess can induce dermatitis, cancer, damage to renal circulation, liver and nervous tissues, while long-term exposure may lead to death [2]. As industrialization train is unstoppable, fight with heavy metal contamination needs innovative remediation strategies. Awareness for treatment and remediation of metal-containing wastes to threshold level before release into natural environment has been growing globally.

Efficient, cost-effective and environment-friendly practices are needed for fine-tuning of waste management. Microbial application is considered an economic and efficient way to remediate HMs from water and soil [3]. Microbes can use inorganic contaminants as a source of energy via activating metabolic processes. Success of the bioremediation process depends on the nature, degree and depth of contaminants, polluted site, environmental policies and cost. Besides, other factors like pH, temperature, nutrient level, oxygen concentrations and abiotic factors also affect bioremediation. The high surface-to-volume ratio of microbes and their potential to remediate metals are considered as the key reason to prefer them.

Studies have demonstrated that indigenous strains that reside in the polluted environments have a significant ability to endure toxic metals. Among the microbial entities, filamentous fungi can grow rapidly and survive in harsh environments. On the contrary, bacteria are ubiquitous biological entities on earth that can reproduce and survive in a variety of environments due to their small size, ease of cultivation on a variety of media and rapid growth [4]. Over time, they have developed resistance against toxic metals to survive in polluted areas because of the high surface-to-volume ratio.

Mostly, metal-resistant fungal strains have been suggested as bioagents for remediating metal-contaminated sites [5]. The composition of fungal cell wall provides active sites for metal sequestration. Preliminary step in the biosorption is a passive process involving various metal binding activities like physical adsorption, ion exchange and complexations, while active process allows the metal to penetrate in the cells. Bacterial application for remediation is a low-input biotechnological practice that is safer and more reliable than conventional methods and can improve soil fertility, characteristics, and quality. For remediation of polluted sites, different resistant bacterial strains have been used for decades [6]. The mechanisms adopted by bacteria to detoxify the metal contaminants are metal exclusion, active transport of metal, biosorption, bioaccumulation, biomineralization, and biotransformation.

Metal-resistant plant growth-promoting microbes are being used to control the metalliferous sites in a productive and eco-friendly way [7]. These microbes not only boost the plant growth process by producing chelators but also reduce the availability of metals to plants. The aim to remediate metals is possible only if the biological entities can resist and tolerate the toxic metals by their physiological and molecular mechanisms [8]. *Trichoderma pseudokoningii*, isolated from TSW, along with AM fungi have been shown to improve growth of *Tagetes patula* in TSW amended soil [9]. Moreover, a synergistic interaction has been observed among *Trichoderma pseudokoningii* and natural or synthetic PGRs to improve growth in pearl millet grown in TSW amended soil [10].

Tannery solid waste represents a metal toxic environmental situation, and it is presumed that it can harbor metal resistant microbes. This is the first comprehensive study of autochthonous microbes from tannery solid waste and their tolerance levels and biosorption efficiency against HMs.

Therefore, this is the first study of its kind with aims:

1. to isolate and characterize the heavy metal resistance of strains of fungi and bacteria against a particular metal or multi-metal environment
2. to observe the most efficient multi-metal resistant strains in synthetic metal solutions
3. to identify the potent strains on a molecular basis for possible future use in bioremediation of metal laden environment.

## Materials and Methods

### 1. Sampling of tannery solid waste

Tannery solid waste samples were gathered from solid waste landfill site of KTWMA, Depalpur Road, Kasur, Pakistan. Twenty-five random samples were collected with at least 10 m distance between every two samples. The material was taken in pre-labeled sterile bags, transported to the laboratory and stored at 4°C for further use.

### 2. Isolation of fungi

One gram of TSW sample from each bag was suspended into 10 mL of sterile water followed by serial dilutions up to 10^6^. About 50 μL dilution was pipetted on 2% ME agar plates using a glass spreader under complete aseptic conditions in a culture room followed by incubation at 25±3°C for 5-7 days. The fungal colonies that appeared were isolated in new plates. Fungal strains were purified as single spore isolates from mature cultures by spreading conidia with a sterile platinum loop.

### 3. Morphological characterization of fungal strains

The fungal strains grown on ME agar medium were morphologically characterized. For slide preparation, a small mycelial plug was mounted onto the slide in lactophenol followed by observation under compound microscope. The morphological characterization up to genus level was performed according to the taxonomic key provided by [11] and [12].

### 4. Isolation of bacteria

One gram of TSW sample was mixed into 10 mL of autoclaved distilled water and serial dilutions (up to 10^6^) were prepared. By using spread plate method, 50 μL sample was pipetted onto LB agar plates followed by incubation at 37°C for 24 h. Based on morphological properties, bacterial colonies were selected and streaked on new agar plates to get single purified colonies.

### 5. Biochemical characterization of bacterial strains

The biochemical characterization of bacterial strains was performed by the methods of [13]. The test performed under biochemical characterization were gram staining, spore staining, catalase, oxidase, starch test, Voges Proskauer, and methyl red.

### 6. Heavy metal resistance assay of fungi

Metal resistance ability of isolated fungal strains was determined by following the protocol of [14]. Pour plate method was selected for screening the resistant fungal strains against Cd, Cr, Cu, Hg, Pb and Zn. The ME agar medium was modified with different metal concentrations ranging from 10 – 1050 mg L^-1^. Fungal disc of 3 mm size with actively growing hyphae was cut aseptically from each isolate and cultured on ME agar plate followed by incubation at 25±3°C for 7 days. After incubation, the growth of fungal strains on metal-containing and control plates was observed. The strains showing resistance at low metal concentration were exposed to higher concentrations and minimum inhibitory concentrations (MICs) of each strain against the metal was determined.

### 7. Heavy metal resistance assay of bacteria

Metal resistance potential of isolated strains was determined by following the method of [15]. Agar well diffusion technique was used for screening of metal-resistant bacterial strains against Cd, Cr, Cu, Hg, Pb and Zn. After incubation at 37°C for 24 h, the zone of inhibition was measured as an indicator of sensitivity. The strains showing resistance at low concentrations were further exposed to high concentrations and their sensitivity was measured.

### 8. Metal biosorption potential of fungal strains

Based on the metal resistance assay, five fungal strains having the maximum metal resistance potential were chosen for biosorption test. Biosorption efficiency of the selected resistant strains was observed at 500 mg L^-1^ concentration (Pb, Cr, and Zn) and 200 mg L^-1^ (Cd and Cu). A disc of ME agar medium having actively growing mycelium of each fungus was cut aseptically and suspended into metal containing-ME broth medium. Flasks were incubated at 25±3°C on an orbital shaker set at 150 rpm for 7-10 days followed by filtration. The supernatant was digested in nitric acid and perchloric acid (3:1 ratio), followed by filtration (Whatman no. 42). The sample was diluted up to 50 mL with distilled water [16]. The total metal concentration was determined on an atomic absorption spectrophotometer and the biosorption efficiency (%) was calculated by the formula:

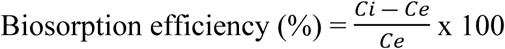

Where, Ci represents initial concentration of metal in the solution and Ce represents final concentration of metal in the solution at equilibrium.

### 9. Metal biosorption potential of resistant bacterial strains

The ten strains having maximum metal resistance were selected and inoculated into LB broth medium followed by incubation at 37°C and 150 rpm until the O.D reached to 0.6 at 600 nm. Metal solution (Cd, Cr, Cu, Pb and Zn) of 500 mg L^-1^ concentration was added into each flask separately including control. Flasks were re-incubated at 37°C for 24 h followed by centrifugation at 5000 rpm for 15-20 min. The supernatant was collected and digested in double volume of concentrated nitric acid on a hot plate at 100°C until the volume reduced to half. The sample was filtered through filter paper (Whatman no. 42) and diluted up to 100 mL using distilled water. The reduction in total metal content was determined on an atomic absorption spectrophotometer and the metal biosorption capacity (%) was calculated following the method of [17].

### 10. Molecular characterization of resistant fungal strains

#### a. DNA extraction

DNA of the five resistant *Trichoderma* strains was extracted using CTAB method [18]. Lyophilized fungal mycelia were homogenized in 2% CTAB extraction buffer followed by incubation at 65°C. After centrifugation, the supernatant was collected in a new microcentrifuge tube and 2 μL of RNase was added into the reaction mixture succeeded by incubation at 37°C for 15 minutes. Next, purification step was carried out by adding an equal volume of phenol: chloroform: isoamyl alcohol (25:24:1) followed by centrifugation at 13000 rpm for 10 minutes. An equal volume of ice-cold isopropanol was mixed with the upper collected aqueous layer followed by incubation at −20°C for 30 minutes. After centrifugation, DNA pellet was washed with 500 μL of 70% ethanol succeeded by centrifugation at 12,000 rpm for 5 minutes. Ethanol was discarded, and DNA pellet was dried and dissolved in 50 μL of TE buffer for further use.

#### b. PCR amplification

To carry out PCR reaction, 50 μL reaction mixture was prepared by adding 2 μL of template DNA (30-35 ng), 10 μL of 5x Phusion buffer, 1 μL of 10 mM dNTPs, 0.5 μL Taq polymerase, and 2.5 μL of 10 mM primer solutions i.e., forward ITS1F (5’ - TCCGTAGGTGAACCTGCGG – 3’) and reverse ITS4R (5’ – TCCTCCGCTTATTGATATGC – 3’) primers were used to amplify the ITS1 and ITS2 region. The PCR conditions were set with an initial denaturation at 98°C for one minute followed by annealing at 50.8°C for one minute and extension at 72°C for one minute. Final extension was performed at 72 °C for 10 minutes. The PCR products were visualized in 1% agarose gel (w/v) having 0.1 μg mL^-1^ SYBR safe and visualized in gel-doc imaging software.

#### c. DNA Sequencing

The PCR products were submitted to the Cornell Institute of Biotechnology for di-deoxy Sanger DNA sequencing and the obtained sequences were subjected to nucleotide BLAST database via NCBI website (http://www.ncbi.nlm.nih.gov) to determine their homology. The sequences were submitted to NCBI GenBank and accession numbers were obtained. For sequence identification variation, all sequences were clustered by using Clustal Omega software (http://www.ebi.ac.uk/).

### 11. Molecular characterization and DNA sequencing of resistant bacterial strains

#### a. DNA extraction

DNA of ten resistant *Bacillus* strains was extracted using Thermo Scientific GeneJet genomic DNA purification kit.

#### b. PCR amplification

To perform PCR, 25 μL reaction mixture was prepared by adding 1 μL of template DNA (25-35 ng), 12.50 μL of Phusion PCR master mix and 1.25 μL of 10μM primer solutions i.e., forward (5’-AGA GTT TGA TCC TGG CTC AG-3’) and reverse primer (5’-GGT TAC CTT GTT ACG ACT T-3’) were used to amplify the 16s rRNA region. The PCR conditions were set with an initial denaturation at 98°C for 30 s followed by 30 cycles (denaturation at 98°C for 10 s, annealing at 56°C for 30 s and extension at 72°C for 30 s). The final extension was performed at 72 °C for 5 min. PCR products were run in 1 % (w/v) agarose gel with 0.1 μg mL^-1^ SYBR safe followed by bands visualization using gel-doc Imaging software.

#### c. DNA Sequencing

The PCR products were submitted to the Cornell Institute of Biotechnology for di-deoxy Sanger DNA sequencing. Percent homology of sequenced strains was checked using the nucleotide blast database through NCBI. The sequences were submitted to NCBI GenBank and accession numbers were obtained. For sequence identification variation, all sequences were clustered by using Clustal Omega software (http://www.ebi.ac.uk/).

## Results

### 1. Morphological characterization of fungal strains

A total of twelve strains of fungi were isolated from TSW and were characterized up to genus level and some to the species level, based on colony morphology and microscopic characteristics (Table 1). Among the isolates, one strain belonged to *Alternaria*, three to *Aspergillus*, two to *Fusarium* and six to *Trichoderma*.

**Table 1.** Colony characteristics of the isolated filamentous fungi

| Strain ID | Color of colony on ME agar medium | Microscopic attributes | Identified genera/ species |
| --- | --- | --- | --- |
| TSWF-1 | black | Hyphae branched, brown, and septate; conidia pale brown, obpyriform, smooth-walled and ovoid with a short cylindrical stalk | *Alternaria alternata* |
| TSWF-2 | green | Conidia unicellular, subglobose and light green in color on tip of phialides | *Trichoderma* sp. |
| TSWF-3 | light to dark green | Hyphae smooth, branched, and hyaline having phialides with unicellular, subglobose and light green colored conidia | *Trichoderma* sp. |
| TSWF-4 | black | Conidiophores long and septate having metulae and phialides | *Aspergillus niger* |
| TSWF-5 | colony appeared green on the periphery and white in the center | Hyphae smooth, hyaline and having unicellular, subglobose, and green colored conidia on phialides tip | *Trichoderma* sp. |
| TSWF-6 | green | Hyphae smooth, branched and hyaline having phialide carried light green, globose to subglobose, and unicellular conidia | *Trichoderma* sp. |
| TSWF-7 | greyish green | Conidiophores colorless, smooth-walled and long, whereas conidia smooth and subglobose | *Aspergillus penicilloides* |
| TSWF-8 | whitish with cottony and flat texture | Macroconidia hyaline, slightly falcate and septate; microconidia cylindrical or ellipsoidal with false heads | *Fusarium oxysporum* |
| TSWF-9 | purplish-white in color | Hyphae branched and erect having unicellular, cylindrical, and ellipsoidal microconidia | *Fusarium* sp. |
| TSWF-10 | greenish-white | Conidiophores branched having phialides; conidia unicellular, thick-walled, light green and subglobose | *Trichoderma* sp. |
| TSWF-11 | greenish-white | Conidia unicellular, subglobose, thick-walled, and light green conidia on phialide | *Trichoderma* sp. |
| TSWF-12 | olive green | Hyphae smooth, septate and hyaline; conidia green in color, thin-walled and subspherical. | *Aspergillus flavus* |

### 2. Biochemical characterization of bacterial strains

A total of twenty-five bacterial colonies were isolated from TSW. Morphological observations revealed that most of the strains to be rod shaped while only five were round (Table 2). Strains TSW-3, TSW-4, TSW-7, TSW-19, and TSW-24 were gram-negative and all others were gram-positive. Except for TSW-6, TSW-7, TSW-10, TSW-18, TSW-20, and TSW-22, all remaining strains were catalase positive. Most of the strains showed positive results for the oxidase test except TSW-3, TSW-4, TSW-7, TSW-13, TSW-19, and TSW-24. Out of 25 bacterial strains, ten strains demonstrated positive results for MR test, while seven strains were found positive for the VP test. Based on the biochemical characteristics, the isolates were identified up to the genus level as *Micrococcus* spp., *Bacillus* spp., *Klebsiella* spp., *Escherichia* sp., *Pseudomonas* spp. and *Streptococcus* spp.

**Table 2.** Biochemical characteristics of isolated bacterial strains

| Isolates | Gram Stain | Spore Stain | Cell Shape | Catalase Test | Oxidase | Starch | MR test | VP Test | Genus Name |
| --- | --- | --- | --- | --- | --- | --- | --- | --- | --- |
| TSW-1 | +ve | +ve | Round | +ve | +ve | +ve | -ve | -ve | *Micrococcus* sp |
| TSW-2 | + ve | + ve | Rod | +ve | +ve | -ve | +ve | -ve | *Bacillus* sp |
| TSW-3 | -ve | -ve | Rod | +ve | -ve | +ve | -ve | +ve | *Klebsiella* sp |
| TSW-4 | -ve | -ve | Rod | +ve | -ve | -ve | +ve | ve | *Escherichia* sp. |
| TSW-5 | +ve | +ve | Rod | +ve | +ve | +ve | +ve | -ve | *Bacillus* sp |
| TSW-6 | +ve | +ve | Rod | -ve | +ve | -ve | +ve | -ve | *Bacillus* sp |
| TSW-7 | -ve | -ve | Round | -ve | -ve | -ve | -ve | -ve | *Micrococcus* sp |
| TSW-8 | +ve | +ve | Rod | +ve | +ve | +ve | -ve | -ve | *Pseudomonas* sp |
| TSW-9 | +ve | +ve | Rod | +ve | +ve | +ve | -ve | +ve | *Bacillus* sp |
| TSW-10 | +ve | +ve | Rod | -ve | +ve | -ve | +ve | -ve | *Bacillus* sp |
| TSW-11 | +ve | +ve | Rod | +ve | +ve | +ve | +ve | +ve | *Bacillus* sp |
| TSW-12 | +ve | +ve | Rod | +ve | +ve | +ve | -ve | +ve | *Pseudomonas* sp |
| TSW-13 | +ve | -ve | Round | +ve | -ve | +ve | -ve | -ve | *Streptococcus* sp. |
| TSW-14 | +ve | +ve | Rod | +ve | +ve | +ve | +ve | -ve | *Bacillus* sp |
| TSW-15 | +ve | +ve | Rod | +ve | +ve | +ve | +ve | -ve | *Bacillus* sp |
| TSW-16 | +ve | +ve | Rod | +ve | +ve | -ve | -ve | +ve | *Pseudomonas* sp |
| TSW-17 | +ve | +ve | Rod | +ve | +ve | +ve | -ve | -ve | *Bacillus* sp |
| TSW-18 | +ve | +ve | Rod | -ve | +ve | -ve | +ve | -ve | *Bacillus* sp |
| TSW-19 | -ve | -ve | Rod | +ve | -ve | +ve | -ve | +ve | *Klebsiella* sp |
| TSW-20 | +ve | +ve | Rod | -ve | +ve | +ve | -ve | -ve | *Bacillus* sp |
| TSW-21 | +ve | +ve | Rod | +ve | +ve | +ve | -ve | -ve | *Pseudomonas* sp |
| TSW-22 | +ve | +ve | Rod | -ve | +ve | +ve | -ve | -ve | *Bacillus* sp |
| TSW-23 | +ve | -ve | Round | +ve | +ve | -ve | -ve | -ve | *Streptococcus* sp. |
| TSW-24 | -ve | -ve | Round | +ve | -ve | -ve | -ve | -ve | *Micrococcus* sp |
| TSW-25 | +ve | +ve | Rod | +ve | +ve | +ve | +ve | +ve | *Bacillus* sp |

+ve: positive, -ve: negative, MR: methyl red, VP: Voges Proskauer

### 3. Heavy metal resistance assay of fungal strains

In the current study, twelve strains of filamentous fungi were isolated from the tannery solid waste. The maximum metal resistance potential of each was studied against six different metals by culturing them on metal-containing ME agar plates. The results showed that among twelve strains, TSWF-6 exhibited maximum resistance for Cd at 400 mg L^-1^; TSWF-4, TSWF-11 for Cr at 800 mg L^-1^; TSWF-2, TSWF-3, TSWF-8, TSWF-11 for Cu at 450 mg L^-1^; TSWF-2, TSWF-6 for Hg at 100 mg L^-1^; TSWF-2, TSWF-5 for Pb at 1050 mg L^-1^ and TSWF-10, TSWF-11 for Zn at 650 mg L^-1^. The MICs of the isolated strains for each metal was determined as shown in the form of a heatmap in Table 3. The resistance of isolated fungi to the studied metals was in the order of Pb*>* Cr> Zn *>* Cu > Cd *>* Hg. The behavior of metal-resistant strains for more than one metal in the form of venn diagram is also shown in Fig. 1

**Table 3.** Minimum inhibitory concentration (MIC) of fungal strains isolated from TSW against six different metals (mg L ^−1^) shown in the form of a heatmap.

| Fungal Isolates | Pb | Cd | Cr | Zn | Cu | Hg | Order of metal resistance |
| --- | --- | --- | --- | --- | --- | --- | --- |
| TSWF-1 | 700 | 250 | 700 | 450 | 350 | 50 | Pb, Cr > Zn > Cu > Cd > Hg |
| TSWF-2 | 1050 | 150 | 350 | 400 | 450 | 100 | Pb > Cu > Zn > Cr > Cd > Hg |
| TSWF-3 | 1000 | 150 | 750 | 150 | 450 | 10 | Pb > Cr > Cu > Zn, Cd > Hg |
| TSWF-4 | 900 | 350 | 800 | 600 | 300 | 50 | Pb > Cr > Zn > Cd > Cu > Hg |
| TSWF-5 | 1050 | 200 | 650 | 500 | 350 | 50 | Pb > Cr > Zn > Cu > Cd > Hg |
| TSWF-6 | 300 | 400 | 250 | 150 | 150 | 100 | Cd > Pb > Cr > Zn, Cu > Hg |
| TSWF-7 | 900 | 250 | 600 | 550 | 250 | 0 | Pb > Cr > Zn > Cu, Cd > Hg |
| TSWF-8 | 800 | 150 | 350 | 500 | 450 | 50 | Pb > Zn > Cu > Cr > Cd > Hg |
| TSWF-9 | 300 | 350 | 400 | 300 | 200 | 10 | Cr > Cd > Zn, Pb > Cu > Hg |
| TSWF-10 | 550 | 350 | 750 | 650 | 150 | 50 | Cr > Zn > Pb > Cd > Cu > Hg |
| TSWF-11 | 300 | 100 | 800 | 650 | 450 | 10 | Cr > Zn > Cu > Pb > Cd > Hg |
| TSWF-12 | 550 | 150 | 250 | 300 | 300 | 0 | Pb > Zn, Cu > Cr > Cd > Hg |

Green color represents the maximum resistance to metal while red represents the lowest resistance to metal by the isolated fungal strain (n=12).

**Fig l.**
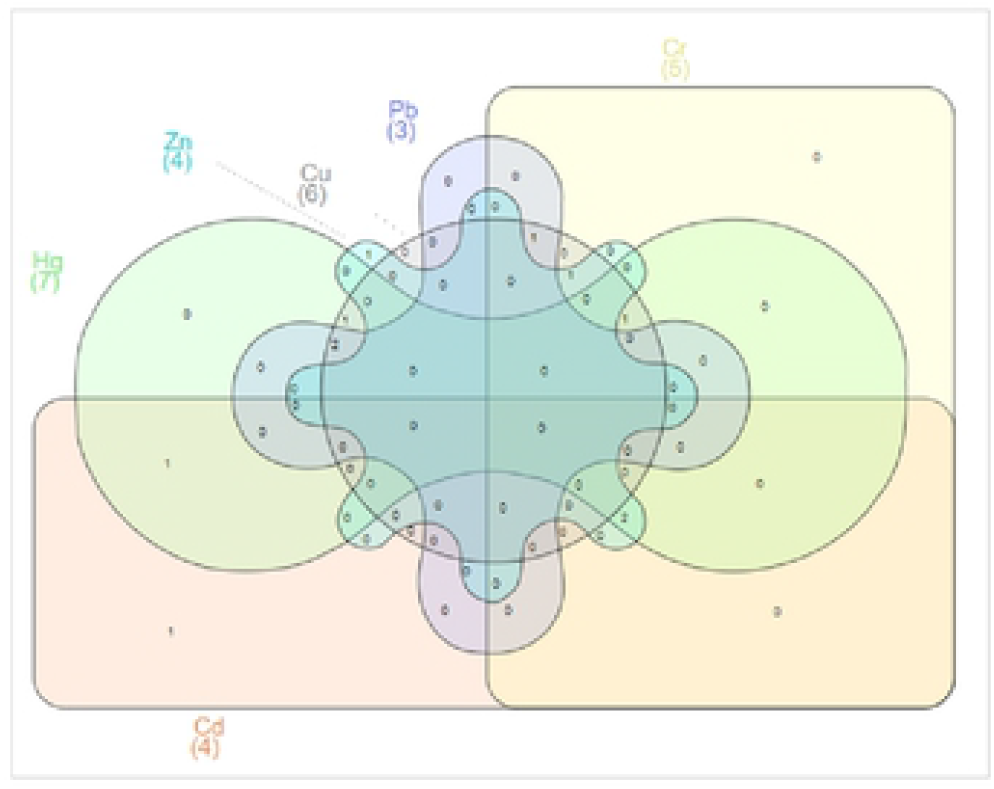
Venn diagram showing the behavior of metal resistant filamentous fungal strains against the heavy metals Cd, Cr, Cu, Hg, Pb and Zn. Among the isolated fungal strains (n=l2), 4 Cd resistant (300-400 mg L^−1^), 5 Cr resistant (700-800 mg L^−1^), 6 Cu resistant (350-450 mg L^−1^), 7 Hg resistant (50-100 mg L^−1^), 3 Pb resistant (950-1050 mg L^−1^) and 4 Zn resistant (550-650 mg L^−1^) strains have been arranged in a Venn diagram.

### 4. Heavy metal resistance assay for bacterial strains

A good resistance against different metals was observed in most of the bacterial strains by zone inhibition plate assay. Diversified results were observed for isolated bacterial strains as shown in the heatmap (Table 4) along with the order of heavy metal resistance against six different metals. The results illustrated that among twenty-five strains, TSW-6, TSW-8, TSW-17 showed maximum resistance for Pb at 1200 mg L^-1^; TSW-2, TSW-14, TSW-15, TSW-21 for Cr at 950 mg L^-1^; TSW-4, TSW-9, TSW-16, TSW-20, TSW-25 for Cu at 650 mg L^-1^; TSW-5, TSW-10 for Cd at 600 mg L^-1^; TSW-4, TSW-19, TSW-22 for Zn at 700 mg L^-1^ and TSW-3, TSW-5, TSW-17, TSW-19, TSW-21, TSW-24 for Hg at 50 mg L^-1^. The behavior of metal resistant strains for more than one metal is depicted in Fig. 2.

**Table 4.** Minimum inhibitory concentration (MIC) of bacterial isolates against six different metals (mg L^−1^)

| Isolates | Pb | Cd | Cr | Zn | Cu | Hg | Order of metal resistance |
| --- | --- | --- | --- | --- | --- | --- | --- |
| TSW-1 | 900 | 150 | 650 | 100 | 300 | 20 | Pb > Cr > Cu > Cd > Zn > Hg |
| TSW-2 | 650 | 150 | 950 | 600 | 600 | 20 | Cr > Pb > Cu, Zn > Cd > Hg |
| TSW-3 | 600 | 100 | 350 | 250 | 600 | 50 | Pb > Cu > Cr > Zn > Cd > Hg |
| TSW-4 | 400 | 50 | 900 | 700 | 650 | 20 | Cr > Zn > Cu > Pb > Cd > Hg |
| TSW-5 | 650 | 600 | 350 | 250 | 250 | 50 | Pb > Cd > Cr > Zn, Cu > Hg |
| TSW-6 | 1200 | 300 | 200 | 250 | 100 | 20 | Pb > Cd > Zn > Cr > Cu > Hg |
| TSW-7 | 850 | 100 | 600 | 100 | 100 | 20 | Pb > Cr > Cu, Zn, Cd > Hg |
| TSW-8 | 1200 | 100 | 650 | 250 | 600 | 20 | Pb > Cr > Cu > Zn > Cd > Hg |
| TSW-9 | 600 | 300 | 200 | 600 | 650 | 20 | Cu > Pb, Zn > Cd > Cr > Hg |
| TSW-10 | 400 | 600 | 600 | 250 | 650 | 20 | Cu > Cd, Cr > Pb > Zn > Hg |
| TSW-11 | 900 | 50 | 200 | 250 | 450 | 20 | Pb > Cu > Zn > Cr > Cd > Hg |
| TSW-12 | 1000 | 150 | 950 | 450 | 600 | 20 | Pb > Cr > Cu > Zn > Cd > Hg |
| TSW-13 | 850 | 100 | 200 | 250 | 300 | 20 | Pb > Cu > Zn > Cr > Cd > Hg |
| TSW-14 | 350 | 150 | 950 | 400 | 450 | 20 | Cr > Cu > Zn > Pb > Cd > Hg |
| TSW-15 | 650 | 100 | 950 | 350 | 300 | 20 | Cr > Pb > Zn > Cu > Cd > Hg |
| TSW-16 | 1100 | 150 | 350 | 400 | 450 | 20 | Pb > Cu > Zn > Cr > Cd > Hg |
| TSW-17 | 1200 | 300 | 350 | 250 | 100 | 50 | Pb > Cr > Cd > Zn > Cu > Hg |
| TSW-18 | 350 | 100 | 650 | 600 | 600 | 20 | Cr > Zn, Cu > Pb > Cd > Hg |
| TSW-19 | 850 | 50 | 350 | 700 | 100 | 50 | Pb > Zn > Cr > Cu > Cd > Hg |
| TSW-20 | 650 | 50 | 600 | 250 | 650 | 20 | Pb, Cu > Cr > Zn > Cd > Hg |
| TSW-21 | 1100 | 150 | 950 | 250 | 100 | 50 | Pb > Cr > Zn > Cd > Cu > Hg |
| TSW-22 | 400 | 300 | 350 | 700 | 300 | 20 | Zn > Pb > Cr > Cu, Cd > Hg |
| TSW-23 | 350 | 50 | 200 | 250 | 450 | 20 | Cu > Pb > Zn > Cr > Cd > Hg |
| TSW-24 | 650 | 100 | 900 | 50 | 600 | 50 | Cr > Pb > Cu > Cd > Zn, Hg |
| TSW-25 | 1000 | 450 | 350 | 250 | 650 | 20 | Pb > Cu > Cd > Cr > Zn > Hg |

Green color represents the maximum resistance to metal, whereas red for the lowest resistance to metal by isolated bacterial strain (n=25).

**Fig 2.**
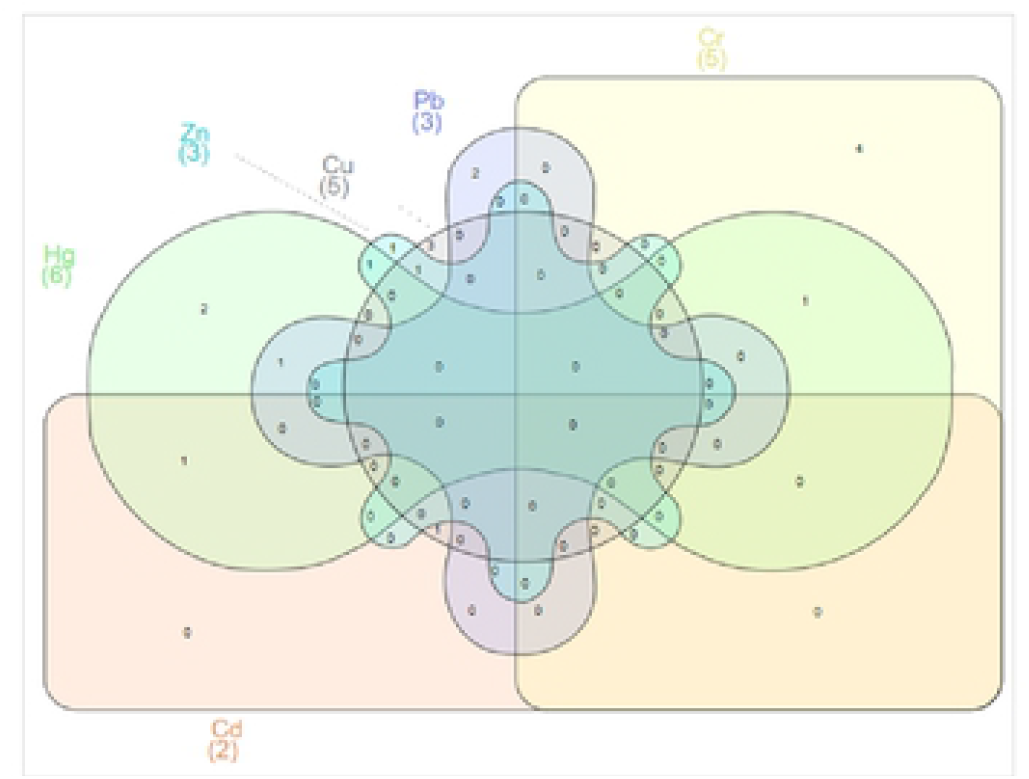
Venn diagram showing the behavior of metal resistant bacterial strains against the heavy metals Cd, Cr, Cu, Hg, Pb and Zn. Among the isolated bacterial strains (n=25), 2 Cd resistant (600 mg L^−1^), 5 Cr resistant (950 mg L^−1^), 5 Cu resistant (650 mg L^−1^), 6 Hg resistant (50 mg L^−1^), 3 Pb resistant (1200 mg L-1) and 3 Zn resistant (700 mg L-^1^) strains have been arranged in a Venn diagram.

### 5. Metal biosorption potential of fungal strains

Biosorption potential of five HM resistant strains against synthetic metal solutions are shown in Fig. 3A. The strain TSWF-10 exhibited 73.7% biosorption for Cr, TSWF-11 exhibited 71.8 % for Cu whereas TSWF-3 showed biosorption potential of 81.7 % for Pb. In the current study, metal resistant strains selected for biosorption showed the maximum removal efficiency for Pb, Cr and Zn compared to Cu and Cd.

**Fig. 3.**
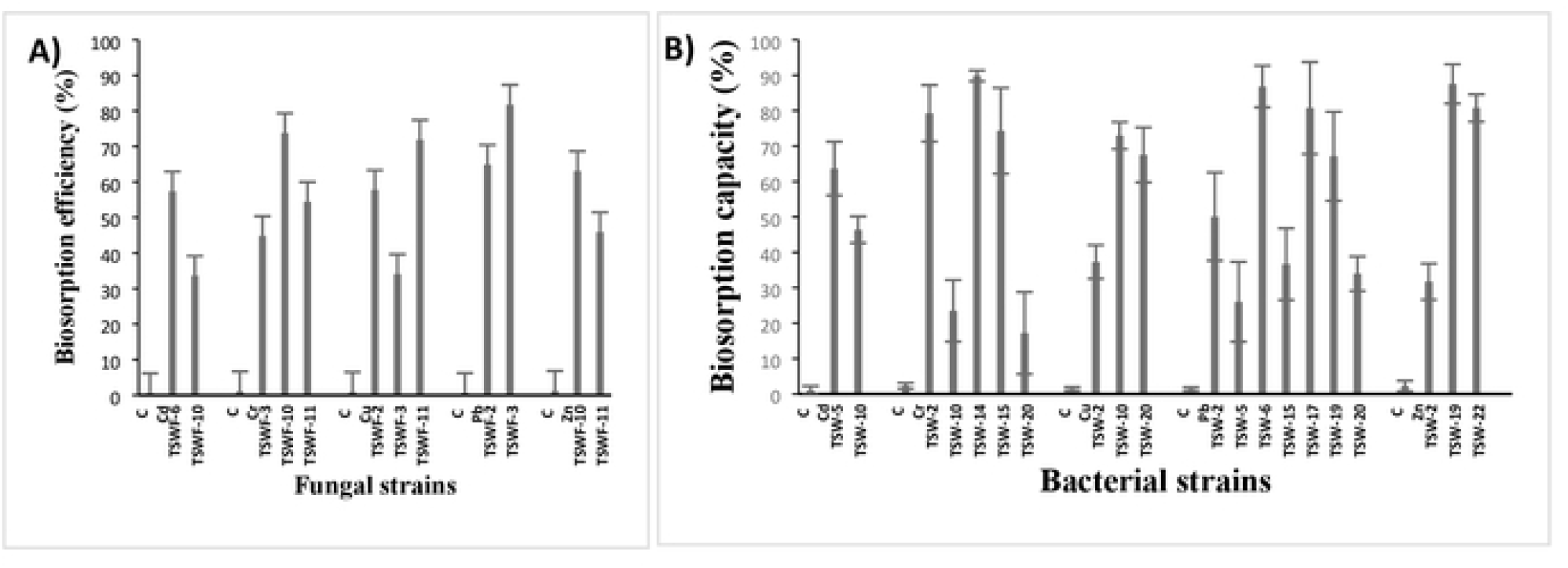
Biosorption efficiency(%) of selected resistant fungal and bacterial strains. **A**. Bar graph of resistant fungal strains against their respective metal concentrations, at 200 mg L^−1^ (Cd and Cu) and 500 mg L^−1^ (Cr, Pb and Zn) concentrations **B**. Bar graph of resistant bacterial strains at 500 mg L^−1^ concentration of metals.

### 6. Metal biosorption potential of bacterial strains

Among the twenty-five strains, TSW-06 and TSW-17 showed maximum biosorption potential for Pb i.e., 86.8 and 80.7 %, whereas, for Cr, 79.2, 89.8 and 74.3 % biosorption potential were exhibited by TSW-02, TSW-14, and TSW-15, respectively. For Zn, 87.5 and 80.7 % biosorption potential was revealed by TSW-19 and TSW-22 (Fig. 3B).

### 7. Molecular characterization of fungal strains

Based on metal resistant assay, five metal resistant *Trichoderma* strains were molecularly characterized by amplifying and sequencing ITS1 and ITS2 regions. The sequence analysis of ITS region revealed that all the five strains belonged to the genus *Trichoderma*. The accession numbers for sequenced *Trichoderma* strains were obtained by NCBI. The statistical analysis of the phylogenetic tree (MEGA Version 10.1.8), generated by bootstrapping (100) and maximum likelihood method showed the similarity index of all the studied sequenced strains with NCBI reported known species as displayed in Fig. 4. The isolated *Trichoderma* spp. were characterized as *Trichoderma lixii* (MW042868.1); *Trichoderma pseudokoningii* (MW042872.1); *Trichoderma pseudokoningii* (MW042876.1); *Trichoderma hamatum* (MW042877.1) and *Trichoderma harzianum* (MW042899.1).

**Fig 4.**
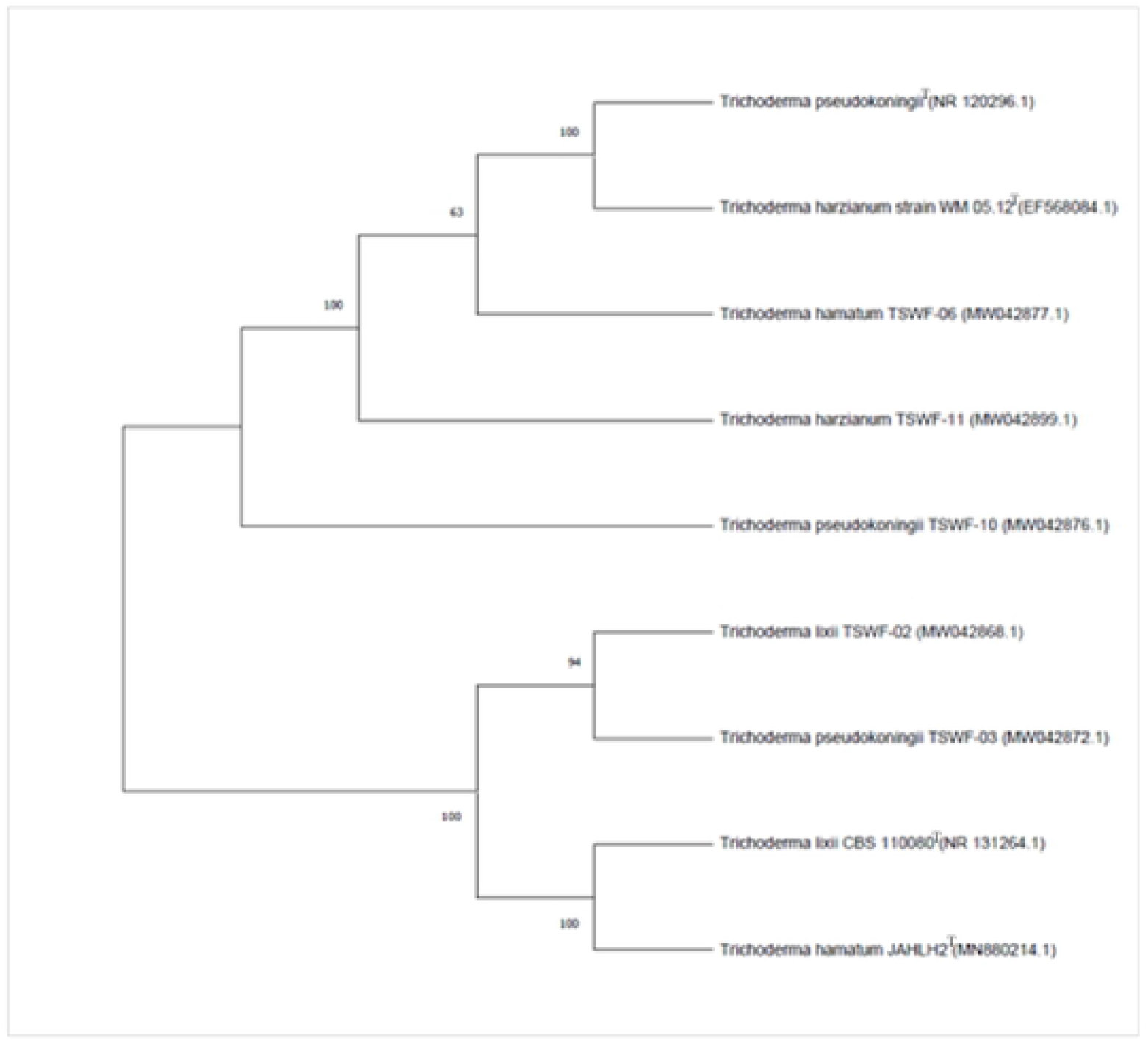
Phylogenetic relationship of five metal resistant *Trichoderma* strains

### 8. Multiple sequence alignments of sequenced fungal strains

Clustal W analysis of DNA sequences of five fungi belonging to the genus *Trichoderma* was executed using bioinformatics tool i.e., Clustal Omega software. The results showed presence of more variation compared to conserved regions as shown in Fig. 5. It was observed that out of the total aligned sequences, 173 base pair long conserved region was observed among DNA sequences of characterized *Trichoderma* strains, symbolized by asterisk. The strains which had more matched base pairs were considered close to each other and vice versa. In the current study, TSWF-3 and TSWF-10 had more matched and less mismatched base pairs and both were identified as *T. pseudokoningii*. Among all strains, more genetic variation was noted in *Trichoderma harzianum* followed by *Trichoderma hamatum* and the variation was represented by ‘grey highlighted area’ in the sequence (Fig. 5).

**Fig. 5.**
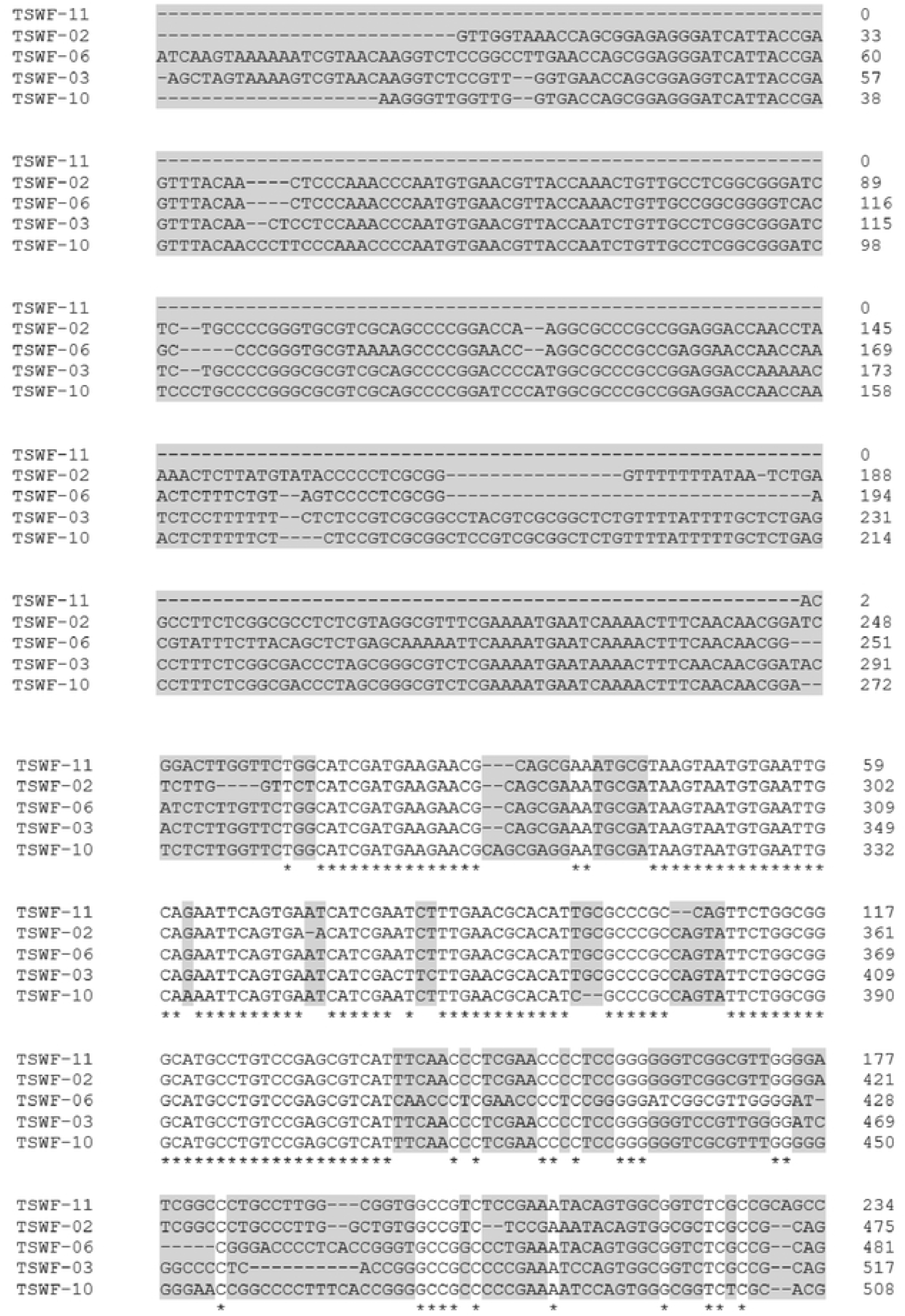
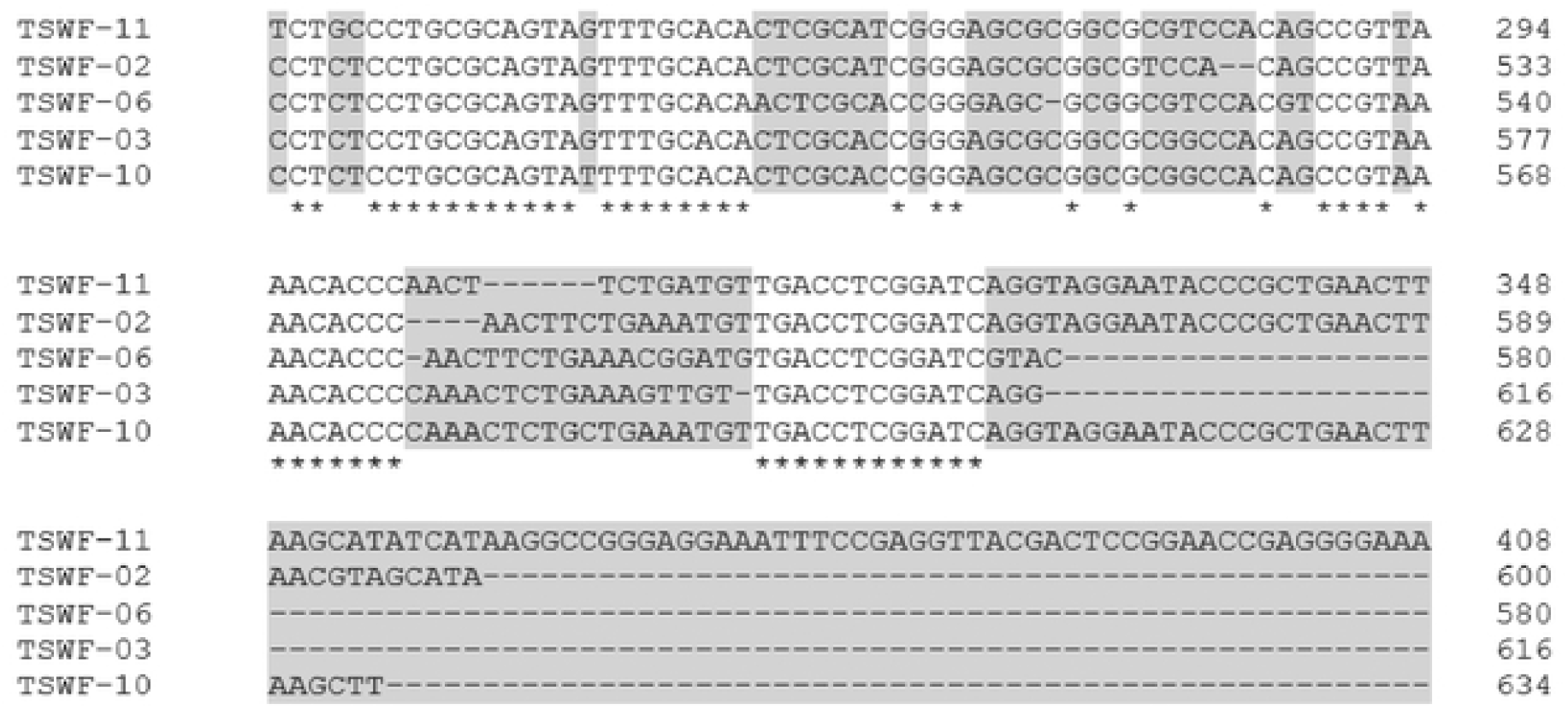
Multiple Sequence Alignment of different metal resistant *Trichoderma* strains showin conserved sequences with asterisks, and genetic variations with grey highlights.

### 9. Molecular characterization of selected bacterial strains

The ten strains of bacteria exhibiting best metal resistance were molecularly identified using 16S rRNA ribotyping technique. The accession numbers for sequenced *Bacillus* strains were obtained by NCBI. Constructed dendrogram results distinguished those selected strains belonging to genus *Bacillus*. Statistical analysis of the constructed phylogenetic tree (MEGA Version 10.1.8), generated by the maximum likelihood method and bootstrapping (100) showed the similarity index of all the selected strains (Fig 6). The *Bacillus* species isolated in this study were identified as *Bacillus xiamenensis* (MT809704.1); *B. velezensis* (MT809705.1); *B. piscis* (MT809706.1); *B. safensis* (MT809709.1); *B. subtilis* (MT810012.1); *B. subtilis* (MT809752.1); *B. subtilis* (MT819963.1); *B. licheniformis* (MT812984.1); *B. cereus* (MT814215.1) and *B. thuringiensis* (MT814283.1). This is the first report of isolation of different *Bacillus* strains from tannery solid waste.

**Fig 6.**
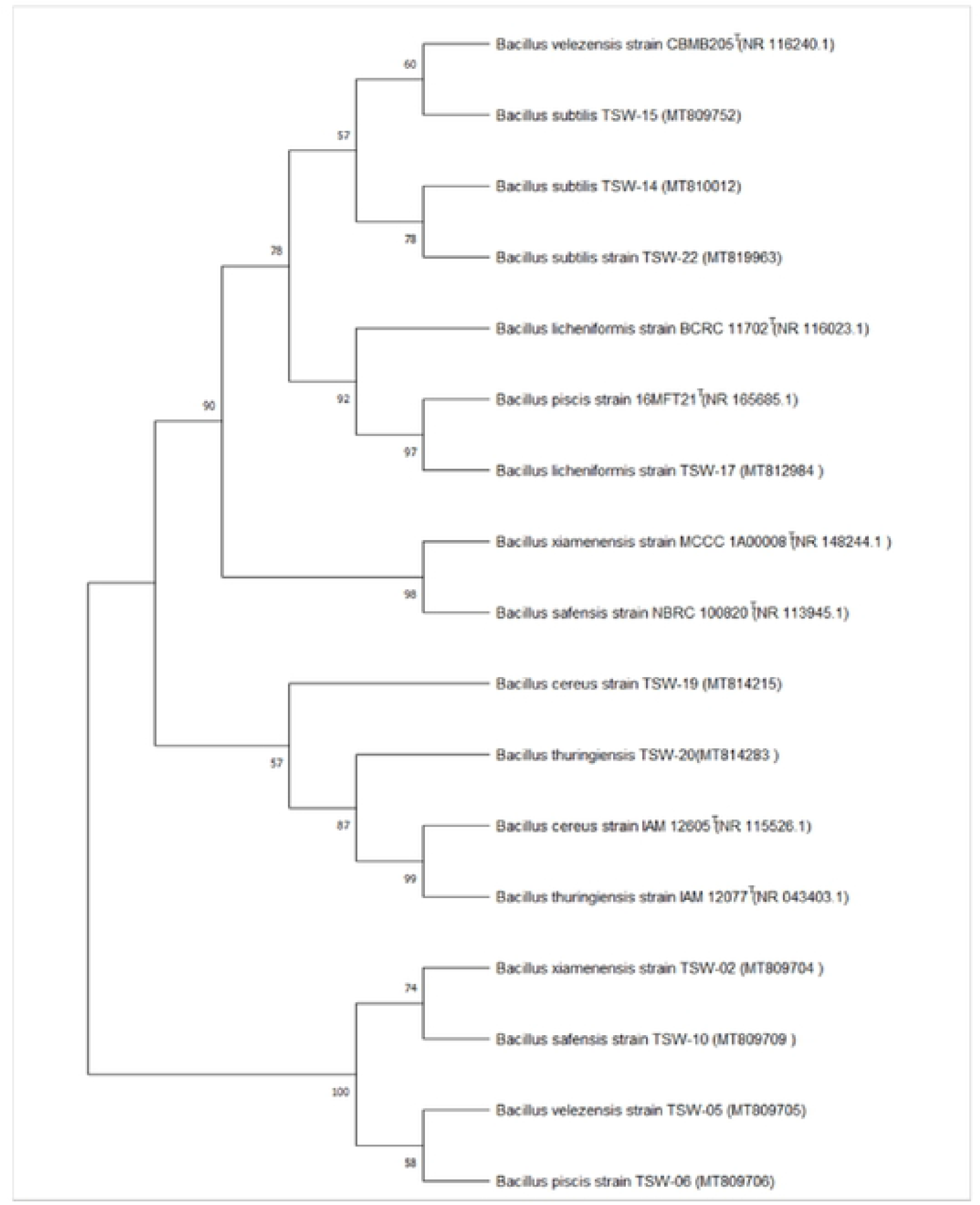
Phylogenetic relationship of ten metal resistant *Bacillus* strains

### 10. Multiple sequence alignments of sequenced bacterial strains

Clustal W analysis of DNA sequences of ten different metal-resistant bacterial strains belonging to the genus *Bacillus* was performed using Clustal Omega software. The results illustrated major ratio of varied region compared to conserved regions as exhibited in Fig. 7. Among the aligned sequences, 90 base pair long conserved region was observed among DNA sequences of characterized *Bacillus* strains, symbolized by asterisk. The strains which had more matched base pairs were deemed close to each other and vice versa. Our finding showed that TSW-14, TSW-15 and TSW-22 had more matched and less mismatched base pairs and all three were named *Bacillus subtilis*. Among all strains, more genetic variation was observed in *B. thuringiensis* accompanied by *B. licheniformis* and the sequence with variation was denoted by ‘grey highlighted area’ in the sequence (Fig. 7).

**Fig. 7.**
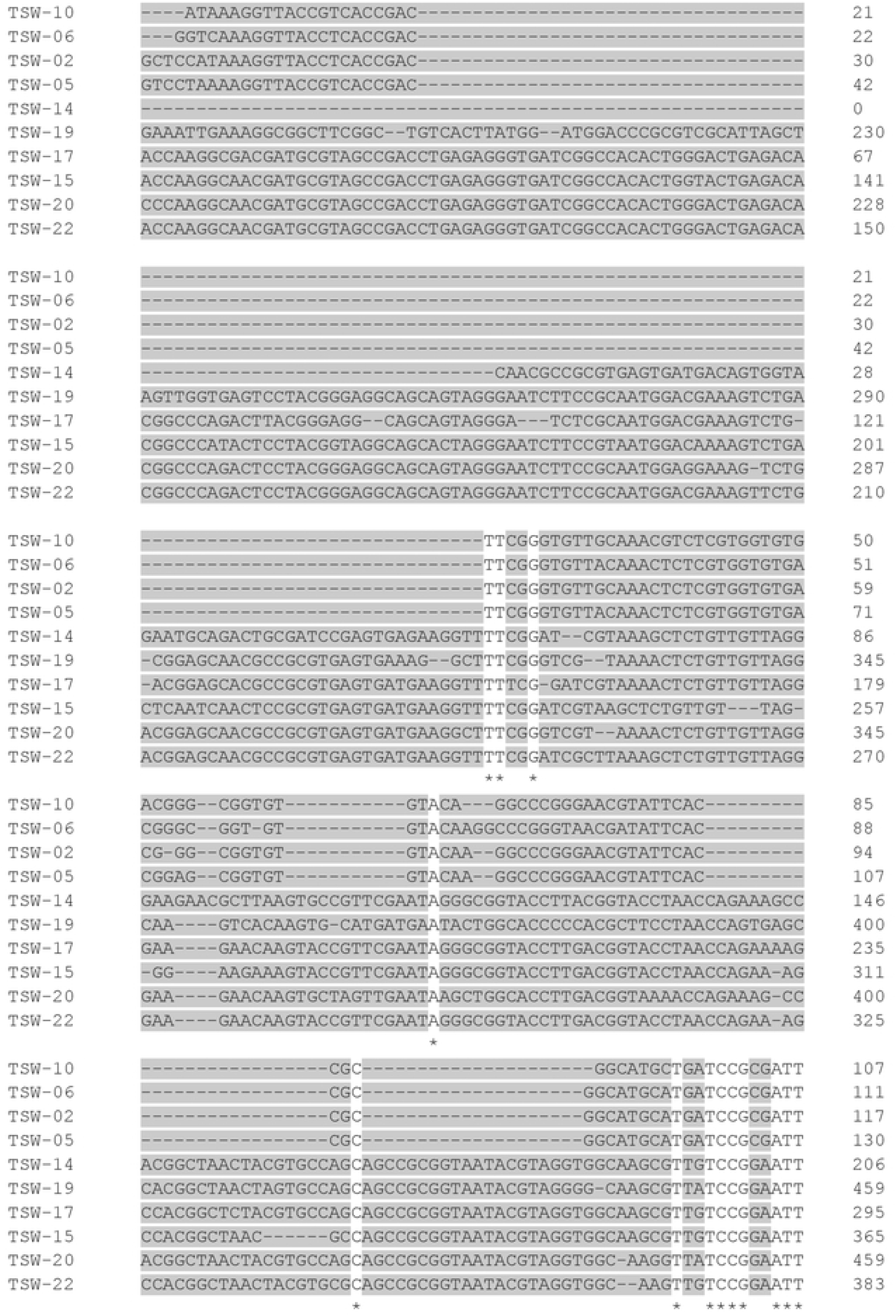
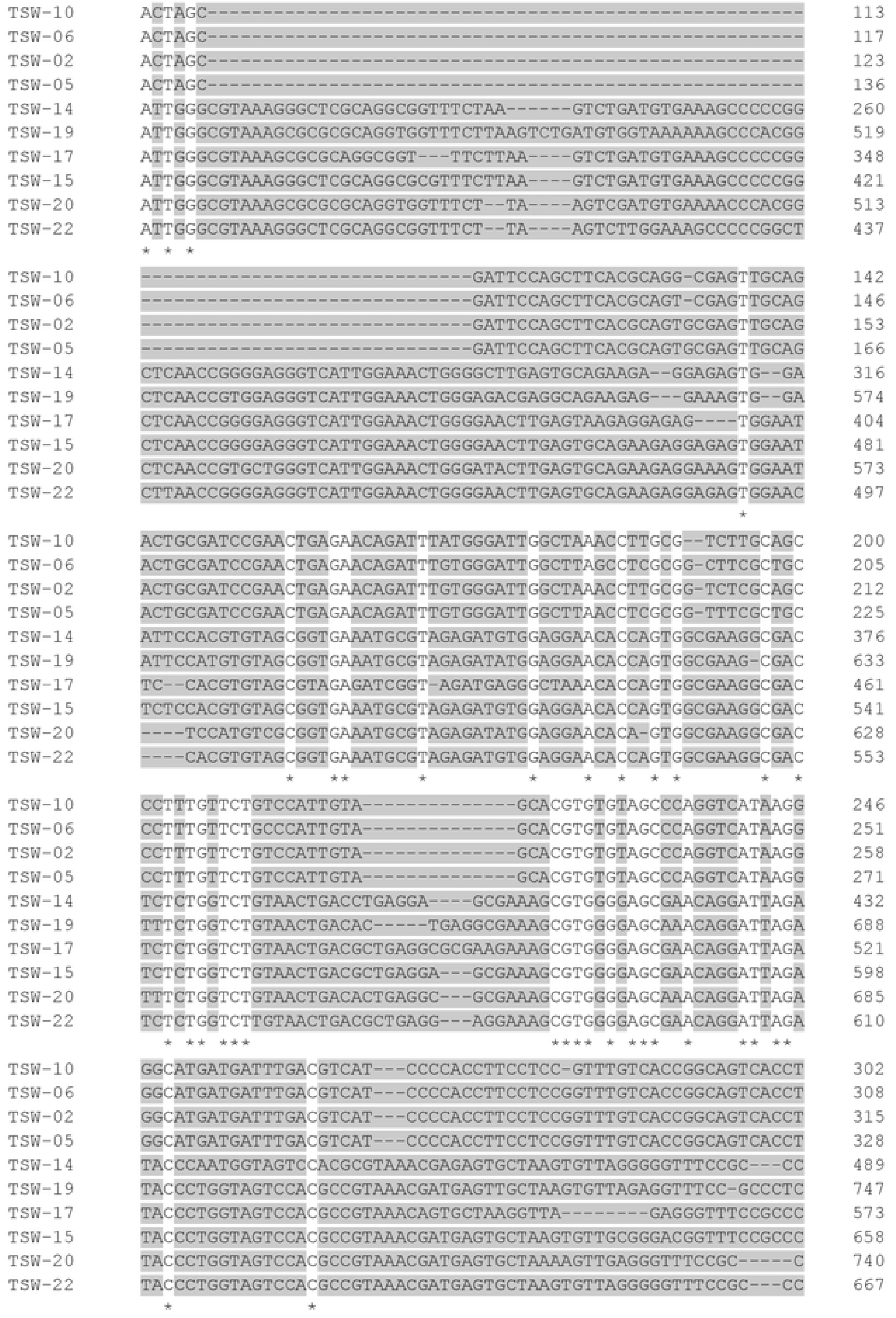
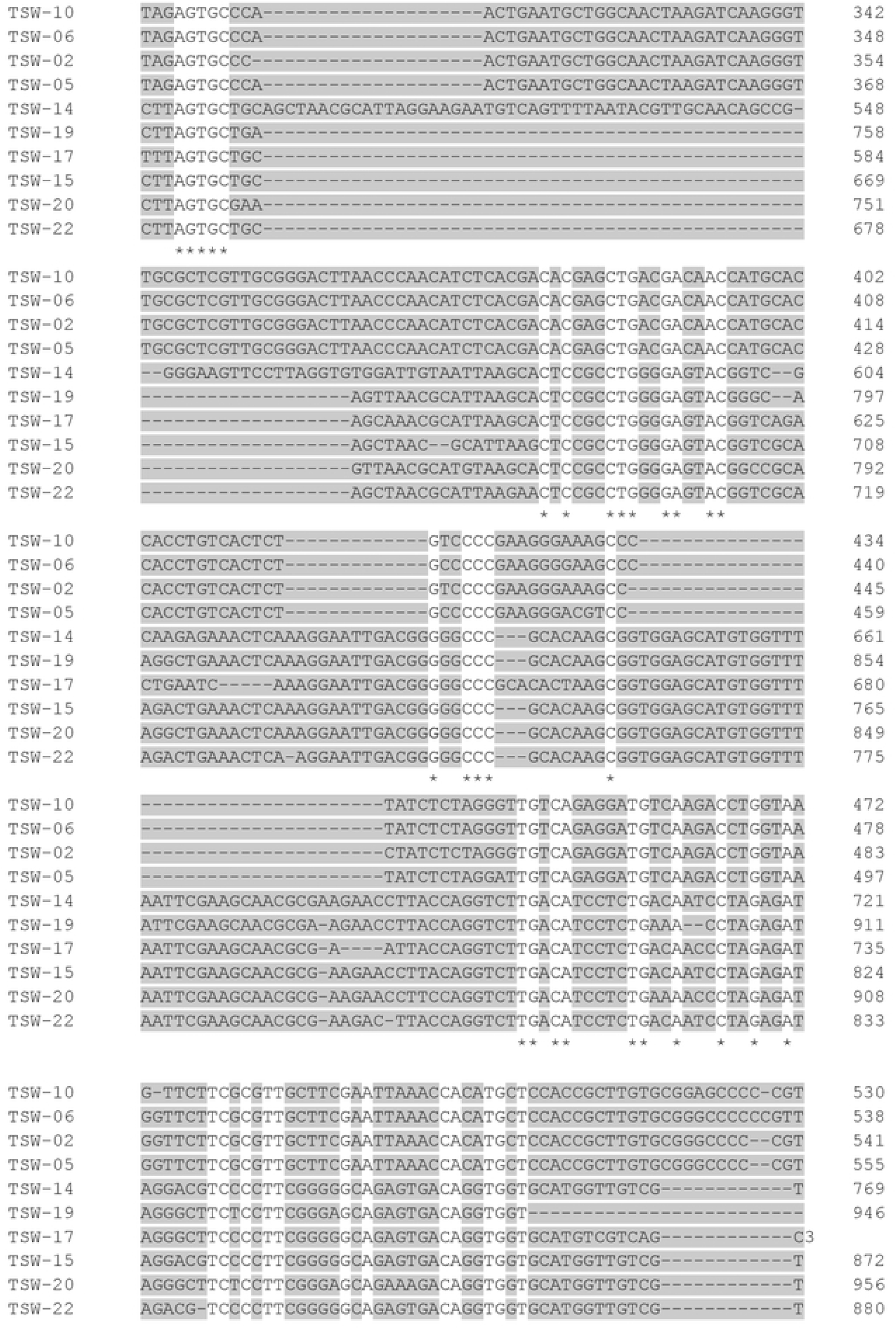
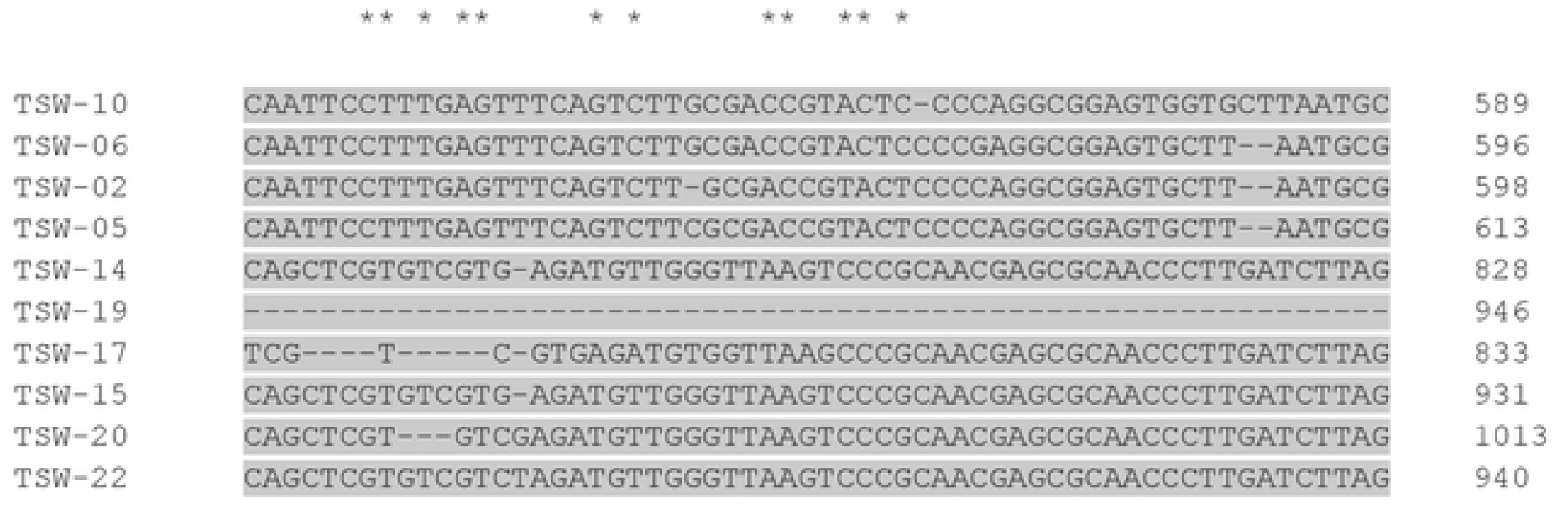
Multiple Sequence Alignment of different metal resistant *Bacillus* strains showing conserved sequences with asterisks, and genetic variations with grey highlights.

## Discussion

Being an industrial city, Kasur faces pollution problems due to the release of metals containing waste from leather industries that pose a threat to the environment, air, soil, human and plants. Tanning industries are considered the major source of metal contaminants, therefore, isolation of metal-resistant strains from such sites may play a vital role in the bioremediation of contaminated sites. Our results showed that among the strains isolated from TSW, one strain belonged to *Alternaria*, two to *Fusarium*, three to *Aspergillus* and six to *Trichoderma. Trichoderma* species have been widely reported from tannery solid waste [9], municipal solid waste [19], metal contaminated soil [20] and mining sites [21]. Numerous studies have shown that *Trichoderma* species resist a high concentration of metals and also improve plant growth under metal stressed environment [22]. The genera of fungi isolated from TSW in the present study have already been observed at metal-contaminated sites. On the contrary, in the current study, biochemical identification results showed that isolated bacterial strains were identified as *Escherichia* (1 spp.), *Streptococcus* (2 spp.), *Pseudomonas* (4 spp.), *Micrococcus* (3 spp.), *Klebsiella* (2 spp.) and *Bacillus* (13 spp.). These results come as no surprise because these genera are the most cultivatable ones. Similar results were demonstrated by [23], who isolated *Pseudomonas* spp., *Bacillus* spp., *and Alcaligens* spp. from leather dye.

In our study, metal resistant potential of twelve fungal isolates against six different metals i.e., Cd, Cr, Cu, Hg, Pb and Zn were assessed, but *Trichoderma* and *Aspergillus niger* showed maximum resistance against the different metals. Several filamentous fungal genera like *Aspergillus, Fusarium, Trichoderma, Humicola* and *Nannizzia* have been reported as metal resistant genera [24]. Increase in metal concentration caused a decrease in fungal growth. The decrease in fungal biomass may appear to be due to high metal concentration and particular MIC of isolated strains. [24] reported that fungal resistance to toxic metals may be attributed to the presence of an effective resistance mechanism. The maximum resistance was observed against Cr and Pb over others. Various resistance mechanisms adopted by fungi for remediation of contaminants include adsorption, oxidation, reduction, methylation, and detoxification. Analogous to our findings [21] reported that *Trichoderma, Rhizopus* and *Fomitopsis* isolated from gold mining sites could resist 400-1000 mg kg^-1^ Pb, Cu, Fe and 25 mg kg^-1^ Cd. [25] reported that *Trichoderma* spp. have the maximum tolerance level against Cd, Cr, Cu, Ni, Pb and Zn. Our study has shown that fungi isolated from TSW have a good resistance against all HMs.

In current study, metal resistant potential of twenty-five bacterial isolates against six different metals i.e., Cd, Cr, Cu, Hg, Pb and Zn were evaluated, but diversified strains of *Bacillus* showed maximum resistance at high concentration of different metals. The bacterial growth at high metal concentrations appears to be due to the presence of an effective resistance mechanism for metal detoxification such as metal efflux, intracellular sequestration, binding with bacterial cell envelopes, and metal reduction. Our findings demonstrated that isolated strains showed maximum resistance potential against Pb and Cr over others. Some of the important bacterial genera used in the remediation of metal-contaminated sites are *Bacillus, Pseudomonas, Acinetobacter* and *Enterococcus* [26]. High metal concentration may lead to a decrease in the growth and biochemical activities of strains, whereas resistant strains have the potential to reproduce at high concentrations.

Fungi and bacteria have a better biosorption potential because of the nature of their cell wall and the functional groups involved in metal binding. Biosorption is described as the elimination of particulates, compounds and metals from a solution by using economical living materials. In the current findings, metal resistant *Trichoderma* strains showed maximum removal efficiency for Pb, Cr and Zn compared to Cu and Cd. Comparable results have been shown by [27], that a high level of Cr could be remediated by applying *Trichoderma* spp. According to [28], among various functional groups, amine and carboxylate groups are important binding sites for metal attachment and biosorption in fungi. However, the success of metal biosorption by fungi depends on the type and concentration of metal, physiological and environmental conditions, availability of nutrients and fungal species [28]. Researchers have reported that fungal strains such as *A. niger, A. terreus, Trichoderma harzianum* and *Rhizopus oryzae* survive in high metal concentrations [29]. Similarly, [30] have shown that *Rhizopus* and *Trichoderma* spp. could resist high concentration of Pb, Cr and Cd. In fact, the presence of *Trichoderma* in highly toxic metal laden environments, already documented by many researchers, has been confirmed in the present study. The fungal strains TSWF-3, TSWF-10 and TSWF-11 showed multi-metal resistance and biosorption and can therefore be employed to clean media contaminated with metals, may be water or soil.

In our study, metal resistant *Bacillus* spp. showed high biosorption for Pb, Cr, and Zn compared to Cu and Cd at 500 mg L^-1^. The reason for high Pb sorption may be attributed to high atomic weight compared to others, which makes it to interact readily with biological components. [31] stated that chromium-resistant strains have the potential to biosorb Cr in the living system either by binding it on the cell wall surface or precipitating it with anions or polymers secreted by bacteria. In the current study, the reason for better biosorption potential of Cr and Zn could be due to the availability of anionic functional groups on bacterial surfaces. The gram-negative strains exhibited better resistance and biosorption for Pb and Zn compared to Cr, Cu and Cd. It is worthwhile to mention that none of the bacterial strains showed multi-metal biosorption, rather individual strains showed affinity for particular metals.

Kingdom fungi is considered the second largest eukaryotic group, ranging from 1.5 - 5.1 million species on earth. Mycologists have encountered the problem of identifying and classifying the wide genera of fungi from a taxonomic perspective. For species-level identification, the ITS regions are considered the fastest and useful part of the rRNA cistron. Over three decades ago, fungal nuclear ribosomal operon primers were used for molecular identification of fungi [32] which helped to generate the sequence of smaller subunit i.e., nrSSU-18S, larger subunit i.e., nrLSU-26S or 28S and Internal Transcribed Spacer (ITS) region i.e., ITS1, 5.8S, ITS2. [33] reported that the maximum likelihood of correct fungal identification could be achieved by sequencing ITS regions. Compared to conventional methods, PCR and Sanger sequencing have been overwhelmingly used for fungal ITS i.e., ITS1, ITS2 and 5.8S. In the current study, five metal resistant *Trichoderma* strains were molecularly characterized by amplifying and sequencing ITS1 and ITS2 regions as given by [34]. The sequence analysis of ITS region revealed that all the five fungal strains belonged to the genus *Trichoderma*. Analogous to our findings, *Trichoderma* species have been widely reported from tannery effluent, municipal solid waste [19], mining sites [21] and metal contaminated soil [20]. Numerous studies reported that *Trichoderma* species have the potential to resist a high concentration of metals and increase plant growth [22]. To best of our knowledge, this is the first report of isolation of different *Trichoderma* strains from tannery solid waste.

Multiple sequence alignment (MSA) is defined as a bioinformatics-based tool for sequence alignment of three or more DNA, RNA, or protein sequences to identify variation. In the current study, maximum intraspecific variation was observed among strains of the same genus i.e., *Trichoderma*. In the current study, more genetic variation was displayed by *T. hamatum* and *T. harzianum*. The possible reasons for these genetic variations could be due to accessibility of metal stress environments to microbes. [35] stated that environmental stress may lead to a high molecular diversity and genetic variability. In the current study, the possible reason for genetic variation seems to be metal stress in the environment.

The ten strains of bacteria exhibiting best metal resistance were molecularly identified using 16S rRNA ribotyping technique. The sequence analysis of the 16S rRNA gene revealed that all the ten strains belong to the genus i.e., *Bacillus*. The *Bacillus* species were widely reported from municipal solid waste [36], municipal wastewater [37], polluted soil [38], coastal environment, tannery effluent [39]. This is the first report of isolation of different *Bacillus* strains from tannery solid waste.

In the current study, multiple sequence alignment results illustrated that *B. subtilis* (TSW-14), *B. subtilis* (TSW-15) and *B. subtilis* (TSW-22) had more matched and less mismatched base pairs. Among all strains, more genetic variation was observed in *B. thuringiensis* followed by *B. licheniformis* and the sequence with variation was represented by ‘grey highlighted area’ in the sequence. The possible reasons for these genetic variations may be attributed to attainability of metal stress environments to microbes.

## Conclusion

Autochthonous microbes isolated from HM polluted environment have the potential to grow and survive in such environment. A variety of filamentous fungi and bacteria isolated from TSW indicated five species of *Trichoderma* and ten of *Bacillus* with a good tolerance and biosorption potential for metals. Molecular characterization indicated the fungi to be *T. hamatum, T. harzianum, T. lixii*, and two strains of *T. psuedokoningii*. On the other hand, the ten bacterial strains were found to be *B. xiamenensis, B. velezensis, B. piscis, B. safensis, B. licheniformis, B. cereus, B. thuringiensis* and three strains of *B. subtilis*. The fungal strains *T. psuedokoningii* (TSW-3, TSW-10) and *T. harzianum* showed a multi-metal tolerance and resistance. Bacteria exhibited metal-specific tolerance and resistance, *Bacillus xiamenensis, B. subtilis* (TSW-14) and *B. subtilis* (TSW-15) against Cr, *B. safensis* against Cu, *B. piscis*^T^ *and B. subtilis* (TSW-17) against Pb and *B. licheniformis* and *B. thuringiensis* against Zn. Thus, the efficient microbial flora can be employed in removing metals from contaminated soil and water.

## Abbreviations

TSW: tannery solid waste
HMs: heavy metals

## Funding

The travel grant was provided by Higher Education Commission (HEC) Pakistan through IRSIP programme to Ms. Hajira Younas (HEC/IRSIP/2020) for visiting Cornell University, Ithaca, New York.

## Acknowledgements

The authors acknowledge the Kasur tannery waste management agency (KTMWA), Depalpur road, Kasur for providing sample for microbial study. We are grateful to HEC Pakistan for providing funding to carry out this research in Cornell University, Ithaca, New York.

## Conflict of interests

The authors declare that they have no conflict of interests.

## Notes

### Competing Interest Statement

The authors have declared no competing interest.

